# Propionate ameliorates diabetes-induced neurological dysfunction through regulating the PI3K/Akt/eNOS signaling pathway

**DOI:** 10.1101/2021.10.15.464493

**Authors:** Qin Wu, Jiajun Dong, Yahong Cheng, Gaofeng Jiang

**Affiliations:** Key Laboratory of Combinatorial Biosynthesis and Drug Discovery, Ministry of Education, Wuhan University School of Pharmaceutical Sciences, Wuhan University, Wuhan, Hubei, P. R. China; Zhejiang University-University of Edinburgh Institute, Zhejiang University, Haining, Zhejiang, P. R. China; Center for Translational Medicine, Tianyou Hospital, Wuhan University of Science and Technology, Wuhan, Hubei, P. R. China

**Keywords:** Propionate, IDDM, Cerebral microcirculation, Neurological dysfunction, PI3K/Akt/eNOS

## Abstract

A large body of research has established diabetes-related cognitive deterioration, sometimes known as “diabetic encephalopathy.” Current evidence supports that oxidative stress, neuronal apoptosis, and cerebral microcirculation weakness are associated with cognition deficits induced by diabetes. The present study explores the effect of propionate on neurological deficits, cerebral blood flow, and oxidative stress in diabetic mice. Propionate can markedly improve neurological function, which is correlated with its capabilities of stimulating nitric oxide (NO) production, increasing cerebral microcirculation, suppressing oxidative stress and reducing neuron loss in the hippocampus. In addition, the results of Western Blotting indicated that the brain-protective function of propionate in streptozocin (STZ)-induced insulin-dependent diabetes mellitus (IDDM) mice is related to phosphoinositide 3-kinase (PI3K)/serine-threonine protein kinase (Akt)/endothelial nitric oxide synthase(eNOS) signaling pathway. In a diabetic mouse model, propionate reduces cerebral microcirculation, hippocampus apoptosis, and neurological impairment. Thus, propionate, now employed as a food preservative, may also help slow diabetes-induced cognitive loss.

**Graphical Abstract:** 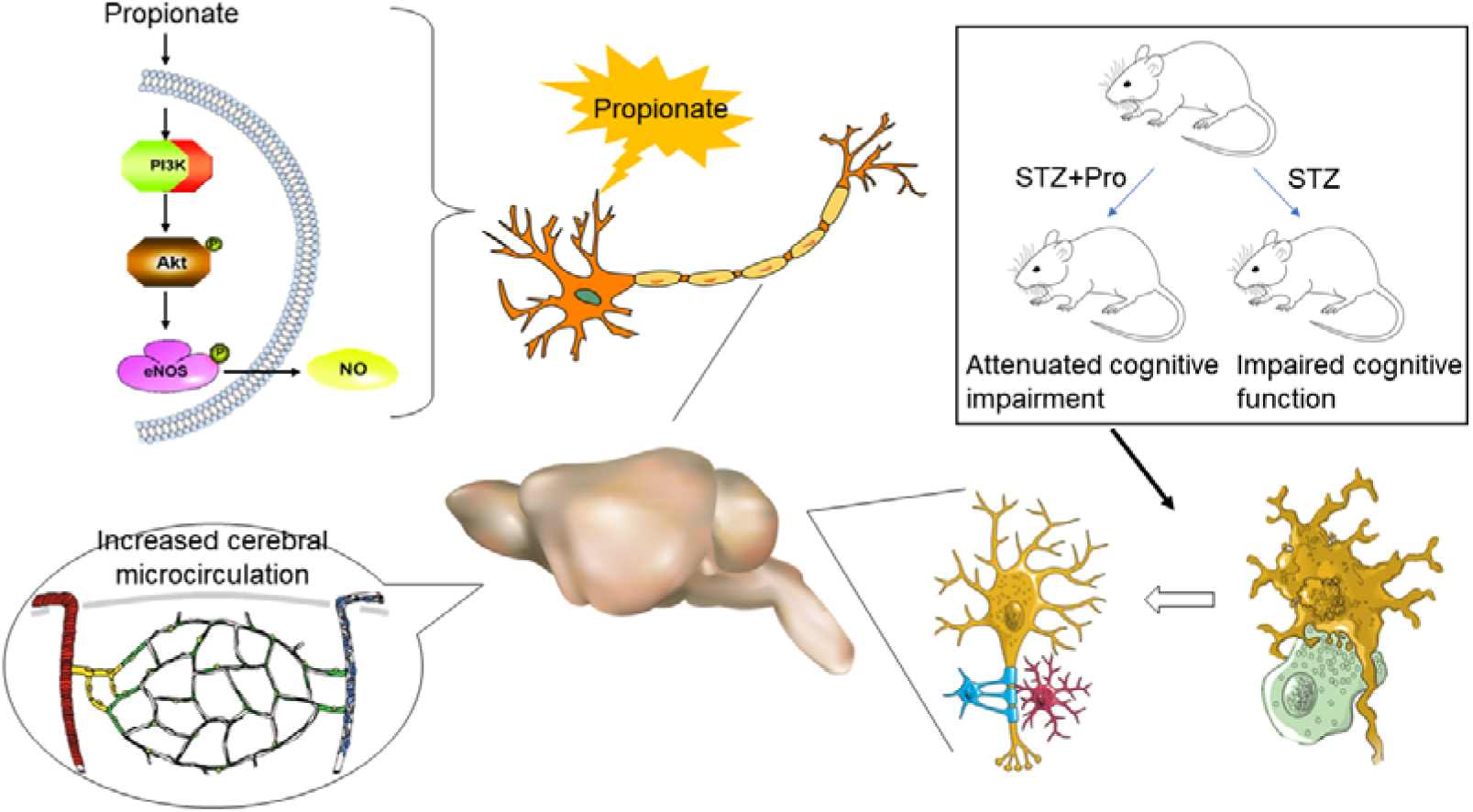

**Highlights:**

- Propionate ameliorates diabetic-induced depression-like behavior, spatial learning and memory deficits.
- Propionate increases cerebral microcirculation by enhancing NO production in diabetic mice.
- Propionate suppresses oxidative stress and reduces neuron loss in the hippocampus.
- The brain-protective function of propionate is related to PI3K/Akt/eNOS signaling pathway.

## 1. Introduction

Cognitive disorder is very common among diabetic patients. Clinical research shows that both type 1 and type 2 diabetes patients are always accompanied by varying degrees of neurological dysfunction ^1, 2^, the incidence of which is related to the course of the disease, the selection of hypoglycemic drugs and other factors ^3^. Slight cognitive impairment may occur in mild cases, and dementia may develop in severe cases ^4^. Therefore, to find effective drugs for the prevention and treatment of diabetic learning nerve function impairment is a research hotspot in this field. Dietary supplementation with nutrients that protect brain tissue from damage may be a promising treatment approach for neurological complications caused by diabetes.

Propionic acid is a kind of short-chain fatty acid (SCFA), which is usually added to cheese, bread, animal feed and artificial spices as a preservative ^5^, and the concentration of propionic acid added as a food additive ranges from 1.0-2.5g/kg. It is reported that propionate plays an important biological role by acting as an antioxidant, anti-inflammatory, antiviral, anti-cancer, etc ^6-8^. Some studies have shown that propionate can inhibit lipid synthesis in rat hepatocytes ^9^. When the concentration of propionic acid was 0.1mmol/L, the synthesis of 50% cholesterol and triglyceride was inhibited ^10^. In addition to reducing cholesterol and fat production, propionate may also be involved in weight control by suppressing appetite ^11^. Propionate enhanced the release of PYY and GLP-1 from human colonic cells considerably in a human-based study. Postprandial plasma PYY and GLP-1 were considerably elevated after ingesting 10 grams of inulin-propionate ester acutely. The enteric release of PYY and GLP-1 creates senses of satiation and satiety in the appetite centers of the brain. Besides, it lessened energy uptake ^11^.

In addition to its role in peripheral nerves, more and more evidence indicates that propionic acid may play an important role in the central nervous system ^11^. Propionic acid was found to enter the brain from the bloodstream in a concentration-dependent manner ^12^. Propionic acid is catalyzed to propionyl coenzyme A (CoA) by thiokinase. Then propionyl CoA reacts with carbon dioxide to generate methylmalonyl CoA under the catalysis of propionyl CoA carboxylase. Finally, methylmalonyl CoA is catalyzed to succinyl CoA by the mutase, thus participating in the tricarboxylic acid cycle and ATP synthesis, which provides energy for the brain ^13^.

However, the therapeutic effect and underlying mechanisms of propionate on treating diabetes-related neurological complications have not been studied. Although the pathologic mechanisms of diabetes-induced nervous system impairment are not fully understood, an increasing number of studies have illustrated that decreased angiogenesis induced by oxidative stress is associated with neurological dysfunction ^14-16^. Oxidative stress is the essential pathologic factor in the early phase ^17^. Oxygen radical damages endothelial cells and induce their apoptosis, which greatly restricts the angiogenesis in brain tissue and limits the blood supply to the brain ^18-20^. It also impacts the secretion of neurotrophic factors by endothelial cells, which exhibit a repairing effect to the function of impaired neurons ^21^. Therefore, it will be a novel and effective therapeutic strategy for treating diabetic neurological complications to prevent oxidative stress injury and thus protect endothelial cells and promote angiogenesis.

Excitatory amino acid toxicity, immune-inflammatory reaction, free radicals and nitric oxide damage are the main factors leading to neuron apoptosis ^22^. Recently, the PI3K/Akt signaling pathway has attracted much attention due to its regulatory role in neuronal apoptosis ^23, 24^. Chemoattractant receptors are known to activate PI3K, a signaling enzyme. When PI3K is activated, it produces the secondary messenger plasma membrane intrinsic protein 3, which interacts with the signaling proteins AKT and phosphoinositide dependent kinase-1 (PDK1). AKT is activated as a result of the combination causing PDK1 to phosphorylate Ser308 in AKT. It is well known that AKT is a significant PI3K effector. ^25^. Phosphorylation of Akt and nitric oxide synthetase is an essential pathway of NO synthesis in endothelial cells ^26-28^. However, whether the intervention of propionate is capable of influencing PI3K/Akt/eNOS signaling pathway is unclear.

The effects of propionate in animal models with diabetic-induced neuronal dysfunction need to be explored. Therefore, the present study focuses on evaluating the effects of propionate on blood glucose level and neuronal function in streptozocin (STZ)-induced diabetic mice. The effects of propionate on cerebral blood flow, oxidative stress, neuronal apoptosis, and PI3K/Akt/eNOS signaling pathway were evaluated at the cellular or protein level through basic pathological tests and biochemical tests enzyme-linked immunosorbent assays, and western blot. This study provides not only a theoretical and experimental foundation on propionate effects of alleviating diabetes-related neurological complications, but also a novel economical, effective and side-effect-free candidate compound for the treatment of diabetic patients with a neurological disorder.

## 2. Materials and methods

### 2.1. Reagents and Chemicals

Sinopharm Chemical Reagents Co., Ltd supplied sodium propionate (> 99 percent purity) (Wuhan, China). Sigma-Aldrich provided the streptozotocin (STZ) and metformin (St. Louis, Mo, USA). STZ was dissolved in citrate buffer, while metformin and sodium propionate were dissolved in distilled water.

### 2.2. Animals

Sixty male Kunming adult mice weighted 18 to 22 grams were used. Every five mice shared one cage and all mice had free access to food and water, under a 12-hour light/12-hour dark condition. Mice were cared strictly compliant with the guidance of the National Institutes of Health Guide for the Care and Use for Laboratory Animals and all animal experiments were performed with the approval of the Institutional Animal Care and Used Committee of Wuhan University (AUP no. S2020110121). Mice were randomly divided into 6 groups, 10 mice for each group. Mice in the normal group (N=10) were not intervened, and 50 successfully modelled mice were randomly divided into five groups of ten mice each: model group, STZ + propionate (37.5 mg/kg), STZ + propionate (75 mg/kg), STZ + propionate (150 mg/kg) and STZ + metformin (100 mg/kg) (positive drug control). The drug intervention groups mentioned above were given high, middle, low dosages of propionate or metformin, respectively. Animals were administered treatments in a randomized, double-blind design, and treatments were randomized across cages to avoid cage level effects.

### 2.3. Diabetes experimentally induced and treatment schedule

STZ (35mg/kg, i.p.) produced in citrate buffer (pH 4.4, 0.1M) was used to develop diabetes, whereas control mice received an identical volume of citrate buffer. After seven days of STZ injection, blood glucose levels were evaluated using blood glucose test strips. Mice with fasting plasma glucose levels greater than 11.1mmol/L were chosen for the subsequent study. Once diabetes was proven, mice were given either sodium propionate (37.5, 75, or 150 mg/kg), metformin (100 mg/kg), or vehicle (0.1 ml/10 g distilled water) orally once daily (9:00 a.m.) for 36 days. Fig. 1 displays the experiment’s timeline.

**Fig. 1.**
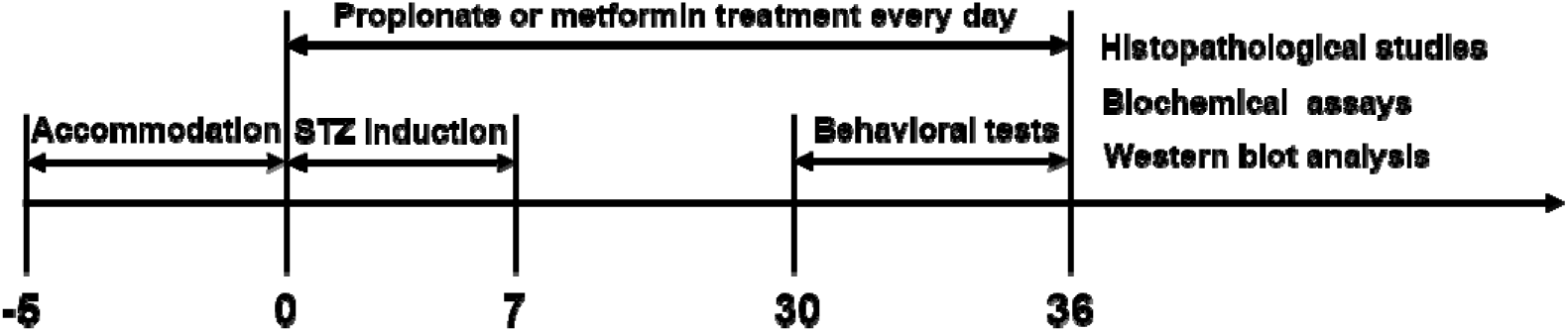
The study design is depicted graphically, encompassing the delivery of drugs, neurobehavioral assessment, histological assessment, biochemical analysis, and protein expression processes.

Propionate levels in foods such as bread range between 1.0 and 2.5 g per kilogram. The dose of propionate used in the present study was determined based on the dose applied as a food preservative. The dosing range for propionate was determined based on our preliminary tests, which demonstrated that propionate was ineffective at doses less than 37.5 mg/kg. However, animals would have difficulty taking propionate at doses more than 75 mg/kg. Thus, we established three dosage groups: high, moderate, and low in order to assess the anti-epileptic effect of the three dosage groups.

### 2.4. Blood glucose and glucose tolerance test

Before the experiment, mice were fasted for 16 hours. Blood was obtained by drawing blood from a vein which is 2 mm from the distal end of the tail and baseline glucose levels were assessed using a glucometer ^29^. Mice were intraperitoneally injected with 100 mg/ml glucose solution in sterile water at 2 g/kg body weight. The blood was taken to measure glucose levels at 0, 30, 60, 120 minutes after the sugar was given ^30^.

### 2.5. Body weight recording

All the experimental mice were weighed at 14:00 to 15:00 once per 3 days throughout the experiment.

### 2.6. Nissl staining

To evaluate the change of neuronal numbers, Nissl staining was performed. Mice were euthanized (N=6 per group) and perfused with 0.9% NaCl solution and 4% paraformaldehyde solution for brain fixation. The entire brain was then isolated and further fixated in paraformaldehyde for more than 24 hours. The brain was embedded in paraffin and sliced into 5 μm thick coronal sections. Paraffin sections were dewaxed and rehydrated before being stained with toluidine blue. Three sections were randomly selected from each mouse, and the stained sections were observed and photographed under a light microscope (Nikon, Tokyo, Japan) in the CA3 region of the hippocampus area. The number of neurons in this region was counted at high magnification (200×)^31^.

### 2.7. Cerebral blood flow measurement

A moorFLPI-2 Full-Field Laser Perfusion Imager (Moor Instruments, Axminster, UK) was utilized to measure local cerebral blood flow. Anesthesia was induced with 4% chloral hydrate, following which mice (N=5 per group) were fixed on a stereotaxic frame plate with a constant temperature of 37□ An incision of skin was made to expose the skull surface, where ultrasound gel was applied. local cerebral blood flow (LCBF) was conducted for 5 min, 25 Hz for sampling frequency and 1 s per frame. The data images were analyzed with moorFLPI2 Full-Field Laser Perfusion Imager Review v5.0 software. For each animal, brain region-of-interest (ROI) was drawn, and ROI flux data was obtained from 19 images per scan. The mean value was used for subsequent analysis. All the analysis was blinded to experimental conditions ^32^.

### 2.8. Determination of oxidative stress-related indicators

#### 2.8.1. Measurement of catalase (CAT), superoxide dismutase (SOD) and malondialdehyde (MDA)

Following completion of the behavioral studies, mice (n = 4) were killed. The tissue was subsequently removed from the brain’s hippocampus region. These tissues (10%, w/v) were immediately homogenized in a freezing potassium phosphate buffer (pH 7.4) and centrifuged at 3000 rpm for 10 minutes at 4 °C. The supernatants obtained were collected and subjected to biochemical analysis.

Protein content was quantified with a commercially available BCA protein assay kit (Solarbio Science and Technology Co., Beijing, China). Malondialdehyde (MDA) level, as a marker of lipid peroxidation, was assessed as contents of thiobarbituric acid-reactive substances (TBARS) by spectrophotometry; the level of superoxide dismutase (SOD) was determined by xanthine oxidase method as the index of the antioxidant system (Nanjing Jiancheng Bioengineering Institute). Based on the operation instruction, catalase (CAT) determination was performed by using a commercial CAT assay kit (Nanjing Jiancheng Bioengineering Institute) ^33^.

#### 2.8.2. Detection of reactive oxygen species (ROS) level

Mouse ROS enzyme-linked immunosorbent assay (ELISA) kit was used to determine ROS concentration. The hippocampal tissues of mice (N=5 per group) were extracted, homogenized, centrifuged, and the supernatant was collected. 40μl supernatant was added to 96-well plate, and reagents were applied according to the manufacturer’s instruction. The absorbance of the mixture at 450 nm was determined by a microplate analyzer. The standard curve was obtained by plotting the concentration of the standard solution with the corresponding absorbance. The sample concentration was determined based on the absorbance of the sample to be tested and expressed as U/mgprot.

### 2.9. Assessment of NO level

NO levels were determined using the supernatant of hippocampal homogenates. NO level was assessed for the purpose of evaluating endothelial cell function in the brain vasculature of diabetic mice. In the body or in an aqueous solution, NO is extremely easy to be oxidized into nitrite ion (NO_2_^-^). Low acidic pH value promotes the reaction of NO_2_^-^ with diazosalt amine sulfonate to form diazo compound, which further couples with naphthalene vinyl diamine to form a product is characterized by an absorption peak of 550 nm. The level of NO can be determined by measuring the absorption value. The NO concentration in brain tissue was determined using a NO assay kit and following the instruction of the manufacture’s (Caozhiyuan Biotechnology Co., Ltd.).

### 2.10. Western blotting

Western blotting was performed following a standard protocol. 20 μg of protein sample from the hippocampus per lane was separated through sodium dodecyl sulfate-polyacrylamide gel electrophoresis (SDS-PAGE). The proteins in the gel were subsequently transferred onto polyvinylidene fluoride (PVDF) membranes and then immunological visualized. Primary antibodies of anti-PI3K (1:1000), anti-p-PI3K (1:5000), anti-Akt (1:1000), and anti-p-Akt (1:5000) were used. β-actin (1:5000) was used as an internal reference. The membrane was incubated overnight at 4□, and the chemiluminescence signal of interest was visualized using an imager (BIO-RAD Gel Doc XR, America) ^34^.

### 2.11. Functional tests

Previous studies indicated that STZ induction resulted in the appearance of behavioral abnormalities such as depression and motor deficits in TIDM mice. In this study, animals were rotated through each test on different days. The test sequence remained constant to guarantee that all animals were subjected to similar test circumstances. All tests were conducted in a blinded fashion, with observers unaware of the treatments administered to mice.

#### 2.11.1. Open field test

A black open-field apparatus (50 × 50 × 40 cm, with 25 equal squares) made of lacrylic box was used to assess the exploratory willingness and spontaneous locomotor activity. The center of the arenas was defined as the central nine squares. Mice (N=10 per group) were placed individually into the center area and allowed 5-min free behave. Animal behavior was recorded using a video tracking system and was analyzed automatically using OFT-100 software. Total distance traveled, the time spent in the center and quiescent, and rearing times were calculated. The apparatus was cleaned with 70% alcohol and dried between trials ^35^.

#### 2.11.2. Tail suspension test

A tail suspension test (TST) was performed to evaluate the depression index of animals by observing the time of immobility and agitation ^36^. The instrument for this test is a grey acrylic box (54 cm height) with an open front and a metallic hook at the top. We use laboratory sticky tape to hang mice (N=10 per group) by the tails from the hook. Their movements were captured for 6 minutes via a camera complete with tracking software. The total immobility period and climbing times were calculated by using the event count mode of RWD software. Longer immobility time was an indicator of depression ^37^.

#### 2.11.3. Forced swimming test

The FST was performed for the purpose of determining behavior related to depression ^38^. Each mouse was placed alone in a vertical plexiglass cylinder (140 mm in diameter, 200 mm high) that contained water at a depth of 150 mm and a temperature of 23-25°C, for a period of 8 min. Recorded for the same period, a blinded experimenter scored each mouse’s immobility time for the last 6 min. Immobility was recognized when mice were floating and moving a minimal amount to keep its head and body from becoming submerged. The researcher remained in the room for the duration of the test, and the water was changed between the introduction of each new mouse to prevent odors (N=10 per group) ^39^.

#### 2.11.4 Elevated plus maze test

The elevated plus-maze test is to investigate the anxiety state of animals by using the contradictory behaviors of exploration and the fear of the high position ^40^. The elevated plus-maze has a pair of open arms (28.5×7cm) and two closed arms (28.5×7×14 cm) they are separated by a central platform (7cm×7cm). The apparatus is 50 cm high. Each mouse was placed on the central platform. The number of entries is recorded, and the time spent in the open arm is recorded over a 5 min time session (N=10 per group).

#### 2.11.5. Morris water maze (MWM) task

MWM was performed to test the spatial learning and memory of the animals ^41^. The basic device of the water maze is a circular pool with a diameter of 1.2 meters and a height of 0.6 meters. The water inside is colored milky white with non-fat milk powder. The water temperature in the pool was maintained at 25±1□.

The pool was evenly divided into four quadrants (called south, east, north, west). An invisible platform with a diameter of 8 cm was hidden 1 cm below the water surface in the target quadrant. The whole test contains 2 phases, navigation test and probe test. During the navigation phase, mice (N=10 per group) were released into the pool from randomly assigned release points, one trial lasted 60 s at most, and would be guided to the platform beyond the time limit and allowed to stay for 15 s. Four trials were run for the MWM navigation phase every day and these were conducted on days 1 to 4. On day 5, the platform was withdrawn from the pool and probing tests were conducted. During the probe phase, the platform was removed from the pool, mice were released for 1 min to swim freely. Tracks of mice in the pool were recorded with the Smart3.0 software during the training and probe phase. There were also 60-second intervals used to record and analyze time spent in the target quadrant (the initial platform quadrant), number of platform crossings, and travel velocity. ^42^.

### 2.12. Statistical analysis

All statistics were analyzed using SPSS statistics version 25, and graphs were generated by Prism 6.0 (GraphPad Software, Inc.). Data for the glucose tolerance test (GTT), body weight and escape latencies in the MWM test were analyzed with a two-way analysis of variance (ANOVA). The other outcomes’ measurement data were compared using one-way analysis of variance (ANOVA). The t-test was used to examine the significance of variable comparisons between groups. Arithmetic mean ± standard deviation (SD) was used to indicate statistical differences, where p less 0.05 was considered statistically significant.

## 3. Results

### 3.1. Reduction of fasting blood glucose and improvement of glucose intolerance by propionate treatment

After 16 hours of fasting, diabetic mice’s fasting blood glucose (FBG) climbed considerably. (p<0.001). However, propionate (75 mg/kg) and metformin (100 mg/kg) treatments significantly decreased FBG (p<0.001) and there was no significant difference among the low dose, middle dose and STZ groups (Fig. 2A). In the intraperitoneal glucose tolerance test (IPGTT), the area under the curve (AUC) of blood glucose was calculated and found to be considerably higher in the STZ group than in the Ctrl group (p<0.001) (Fig. 2B-C; Table 1). Comparing with STZ administrated group, propionate administration reduced the AUC (p<0.05 in 37.5 and 150 mg/kg groups; p<0.001 in 75 mg/kg group), and treatment of 75 mg/kg propionate has significant advantages over 37.5 mg/kg and 150 mg/kg of propionate gavage groups. The data clearly showed administration of propionate at a dose of 75 mg/kg effectively decreased the FBG and improved glucose intolerance in IDDM mice. In the remaining experiments, propionate at a dose of 75 mg/kg was selected for intervention.

**Table 1.**
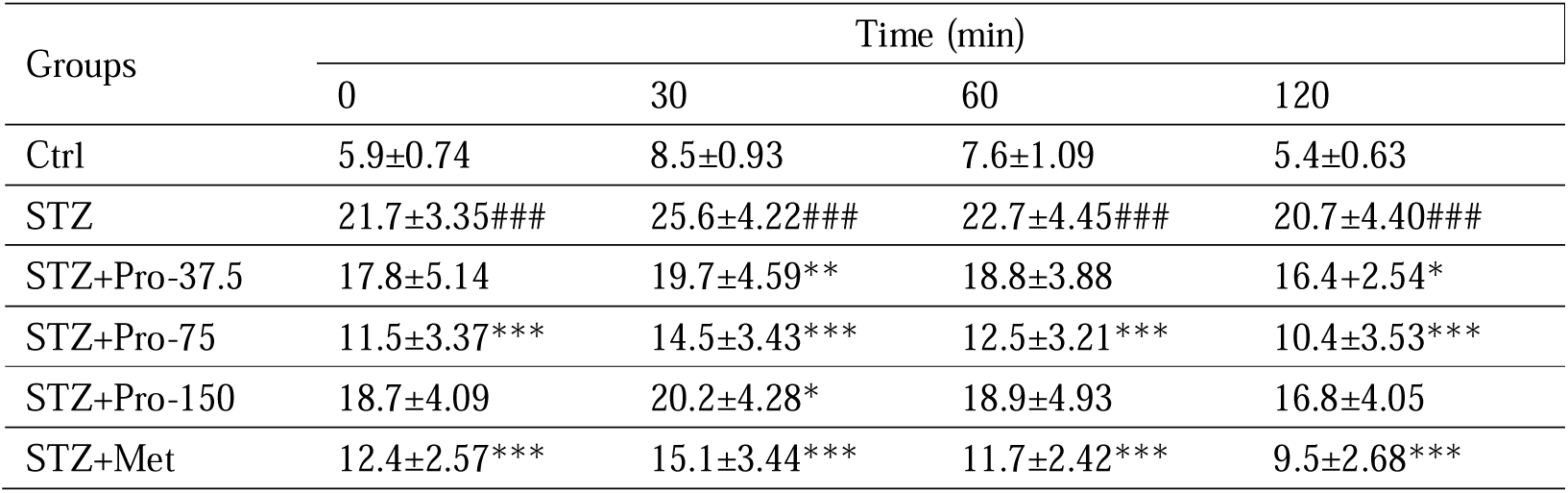
Blood glucose in the IPGTT test. Data was expressed as mean ± SD (N = 10). ###p < 0.001 vs. Ctrl group, *p < 0.05, **p < 0.01, and ***p < 0.001 vs. STZ group. The two-way ANOVA for IPGTT indicated a significant group-by-min interaction.

**Fig. 2.**
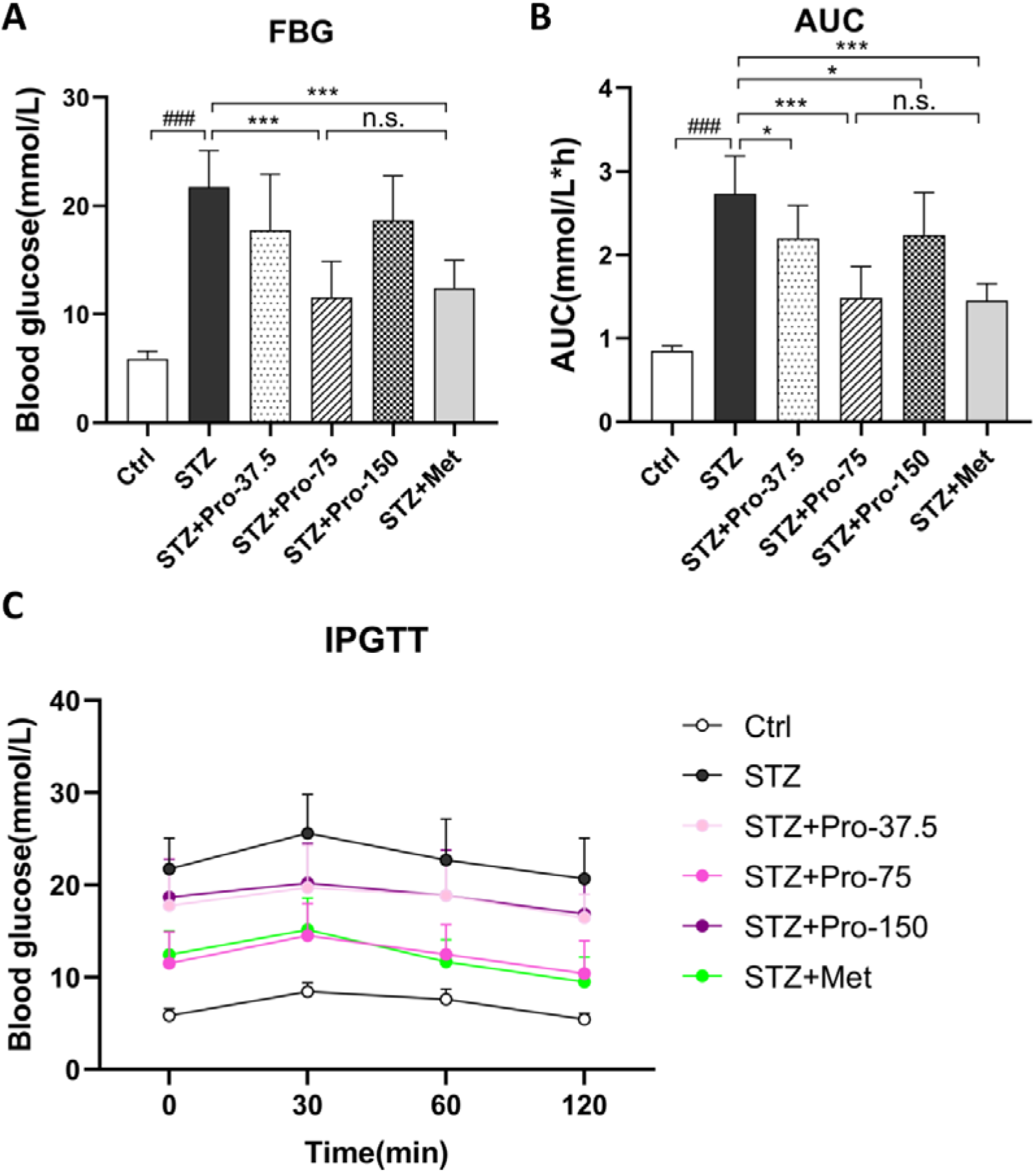
Effects of propionate on FBG and IPGTT. (A) FBG levels; (B) The AUC of blood glucose in the IPGTT test; (C) IPGTT test 36 days after STZ induction. The data are expressed as the mean ± SD (N = 10). ^###^p < 0.001 vs Ctrl group; *p < 0.05, **p < 0.01 and ***p < 0.001 vs STZ group. Data for the IPGTT were analyzed with a two-way analysis of variance (ANOVA).

### 3.2. Propionate ameliorated STZ-induced body weight loss

The control group exhibited a gradual increase in body weight from 19.9±2.50g to 43.8±3.25g. (Fig. 3). Meanwhile, the streptozotocin-induced mice exhibited a significant decrease in body weight from 20.3±3.27g to 33.1±4.28g (p<0.001) in comparison to the former group. Conversely, when compared to the model group, the animals who received propionate (p<0.001) or metformin (p<0.01) exhibited a significant enhancement in their body weight, from 20.3±2.81g to 40.2±2.94g and from 19.2±1.28g to 37.4±1.62g, respectively. The results showed that streptozotocin significantly reduced body weight in the animals compared to the control group, while propionate therapy increased body weight, reversing the weight loss caused by streptozotocin.

**Fig. 3.**
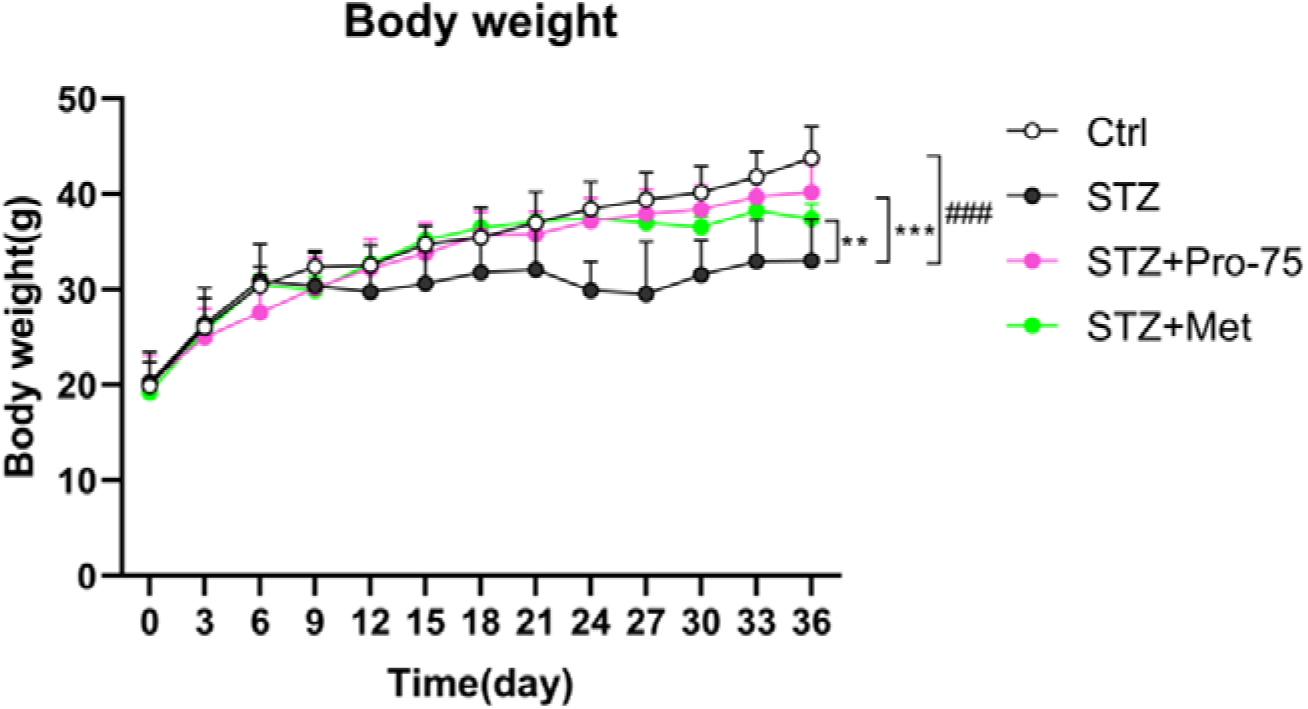
The body weight data of the untreated control group, the streptozotocin exposure group, and the groups treated with propionate and metformin. All data were expressed as mean ± SD (N = 10). ^###^p < 0.001 vs Ctrl group; **p < 0.01 and ***p < 0.001 vs STZ group. Data for the body weight were analyzed with a two-way analysis of variance (ANOVA).

### 3.3. Improvement of local cerebral blood flow (LCBF) by propionate intervention

We determined the effects of propionate and metformin on the LCBF of mice since it can provide objective indicators for cerebral circulation or obstruction. LCBF was measured by Doppler flowmetry (Fig. 4). The STZ-administrated group exhibited a compromised LCBF level with approximately 35.11% decrement (p<0.001). The pre-treatment of propionate significantly increased LCBF comparing with the model group (p<0.01), while metformin treatment failed to reverse LCBF decrement caused by STZ. These observations indicated that propionate could rescue decreased LCBF levels induced by diabetes.

**Fig. 4.**
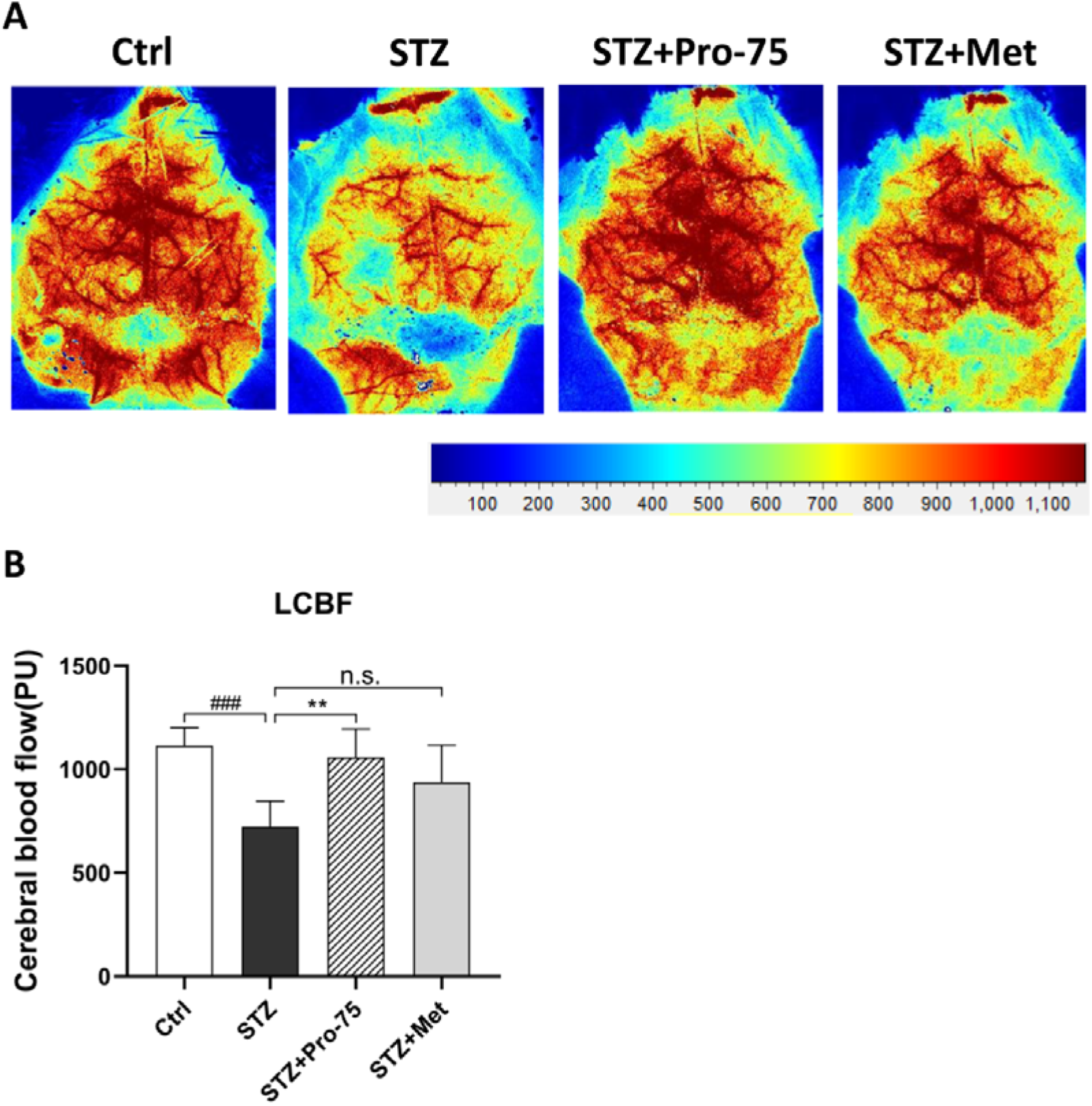
The effect of propionate on LCBF in mice cerebral cortex. (A) Representative images of LCBF in different groups. The magnitude of LCBF is represented by different colors, with blue to red indicating low to high. (B) Quantitative analysis of LCBF in all groups. Data were expressed as mean ± SD (n = 6). ###p < 0.001 vs. Ctrl group, **p < 0.01 vs. STZ group, n.s., no significant difference.

### 3.4. Oxidative stress partially ameliorated by pretreated propionate

#### 3.4.1. Propionate enhanced CAT and SOD levels

The effect of propionate on antioxidase was further studied by measuring CAT and SOD activities. As figure 5A showed, CAT activity was significantly reduced after STZ intervention when compared to the non-treated group (p<0.01). While CAT activity was significantly increased after propionate treatments compared with the STZ group (p<0.01). Furthermore, CAT in the propionate treated group was found to be higher than in metformin-treated group (p<0.05). In addition, SOD activity was obviously decreased in STZ-exposed mice in comparison to the control group (p<0.01). The activity of SOD was restored in mice following propionate or metformin treatment (p<0.05), and no substantial difference between the metformin and propionate groups was found (Fig. 5B).

**Fig. 5.**
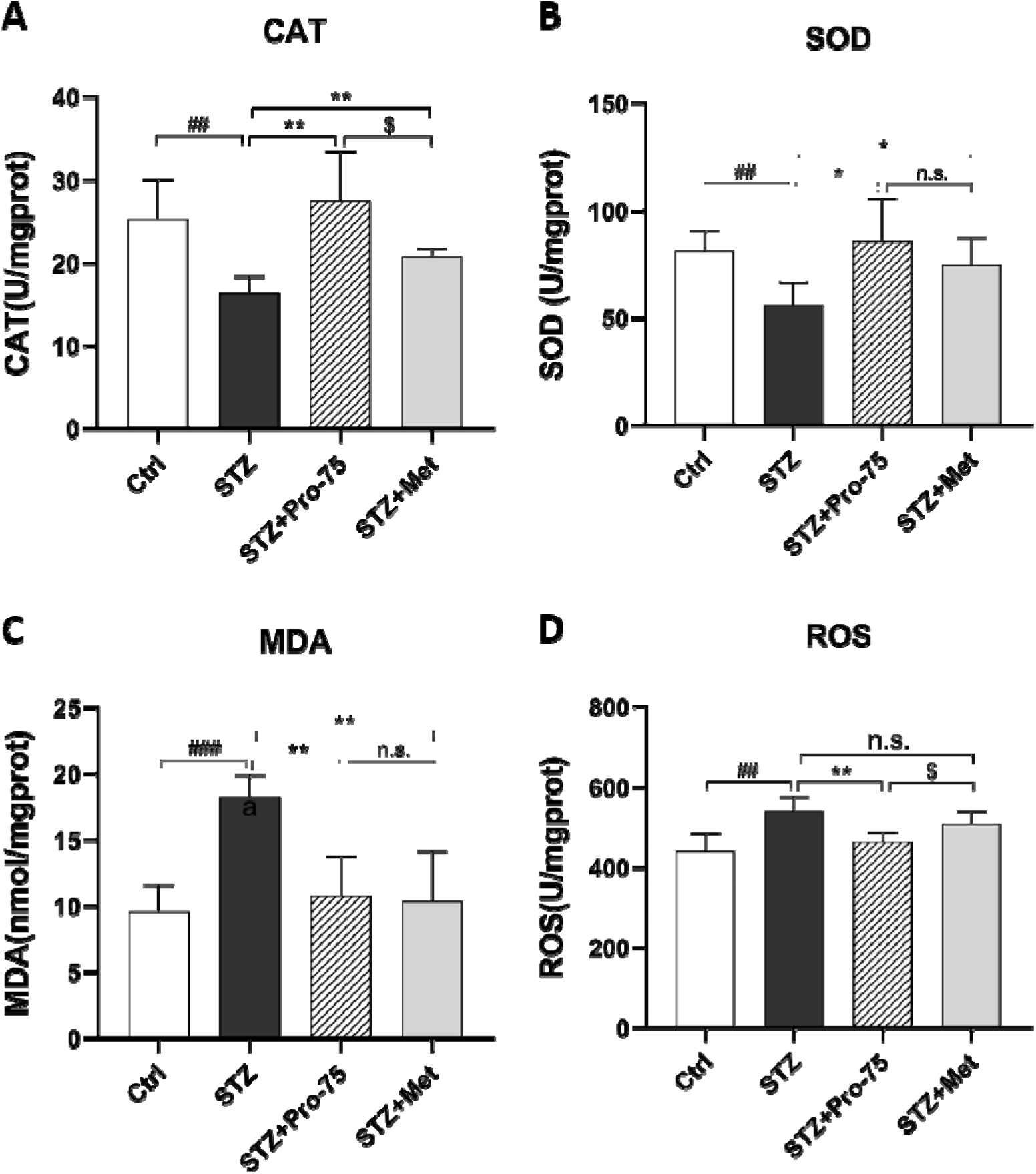
Levels of oxidative stress. (A and B) STZ decreased CAT and SOD. Pretreatment with propionate attenuated these decrements. (C and D) STZ increased MDA and ROS levels in mice brains. Pretreatment with propionate attenuated these increments. Data were expressed as mean ± SD (n = 6). ###p < 0.001 and ##p < 0.01 vs. Ctrl group, *p < 0.05 and **p < 0.01 vs. STZ group, ^$^p < 0.05 vs. STZ + Met group, n.s., no significant difference.

#### 3.4.2. Inhibition of MDA and reactive oxygen species (ROS) accumulation by propionate

To ascertained the potential anti-oxidative effect of propionate, we also measured MDA (a marker of lipid peroxidation) and ROS content in mice hippocampal tissue. The MDA contents in mice hippocampus are shown in Fig. 5C. MDA formation was detected to be significantly elevated in the STZ group compared to the control group (p<0.001). In contrast, significant decreases in MDA levels were found in propionate or metformin treatment groups comparing to the STZ group (p<0.01). No statistical difference was presented between propionate and metformin groups.

The effects of increased production of ROS involve brain damage. ROS can produce membrane damage, change inner proteins, induce lipids denaturation, and damage DNA. ROS concentration was detected in mice brain tissues. Mice administrated by STZ showed significantly higher ROS levels than the control group (p<0.01) (Fig. 5D). ROS concentration was remarkably reversed by propionate intaking compared with the STZ group (p<0.01). While metformin treatments showed little benefit effects. Propionate application had a distinct advantage over metformin administration in reducing ROS concentration (p<0.05). In summary, propionate showed positive effects in scavenging excessive reactive oxygen levels in mice brains.

### 3.5. Propionate enhanced ATP levels

The model group’s ATP level was substantially lower than the control group’s (p < 0.05). It was significantly elevated following propionate pretreatment in comparison to the STZ-induced group. However, metformin had no discernible effect on the level of ATP (Fig. 6A).

**Fig. 6.**
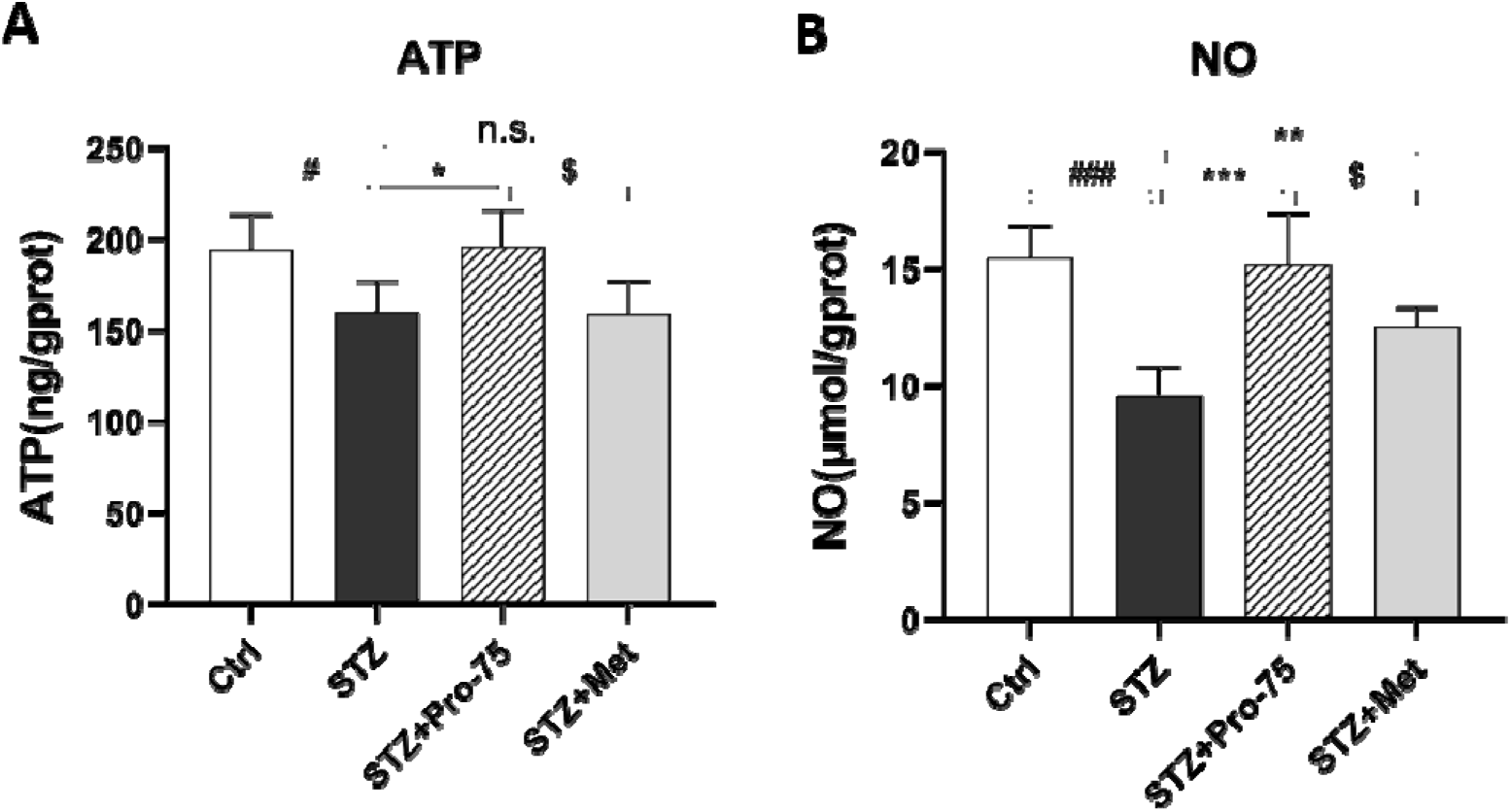
Effects of propionate on ATP level and NO release in brain tissues. Data were expressed as mean ± SD (N = 6). #p < 0.05 and ###p < 0.001 vs. Ctrl group, *p < 0.05, **p < 0.01 and ***p < 0.001 vs. STZ group, ^$^p < 0.05 vs. STZ + Met group. n.s., no significant difference.

### 3.6. Enhancement of nitric oxide (NO) release by propionate therapy

We examed brain NO level to evaluate NO production in brain vasculature. NO concentration was significantly lower in the STZ group (p<0.001): less than two-thirds that of the control group (Fig. 6B). Pre-treatment of propionate or metformin to the STZ-exposed mice raised NO levels by 59.2% and 31.5%, respectively compared with the STZ group (p<0.001 and p<0.01, respectively). The NO level in the propionate group was also elevated in comparison with that in the metformin group (p<0.05). The increased NO concentration indicated an effect of propionate to promote angiogenesis in the brain, with a significantly higher efficacy than metformin.

Propionate can significantly increase the activity of antioxidant enzymes and reduce the level of malonaldehyde as an antioxidant. Although NO is also a product from oxidative stress, it has protective effect on brain in this study. The experimental results showed that the main role of NO is to dilate blood vessels, protect vascular endothelium and increase blood supply to the brain. It can be seen that the protective effect of NO is greater than its oxidative damage. The experimental results obtained and presented are reasonable.

### 3.6. Administration of propionate rescues STZ-induced neuronal loss in the hippocampus

Thirty-six days after STZ administration, we utilized Nissl staining to evaluate the structures of the hippocampus in each group through comparing the number of neurons in optical fields of the CA3 regions across different groups. STZ administration induced obvious lesions in neurons in the CA3 and caused significant neuronal degeneration (p<0.001). On the contrary, metformin and propionate rescued the degradation as demonstrated by the results that numbers of neurons in STZ + propionate (Pro) and STZ + metformin (Met) groups are significantly higher than that in the STZ group (p<0.001). Noteworthily, propionate exhibited a superior effect than metformin (p<0.01). The data indicated that pre-treatment of propionate could persistently alleviate neuronal necrosis resulted from STZ-induction.

### 3.7. Improvement of exploratory and locomotor activity following propionate administration

It is well documented that exploratory willingness and spontaneous locomotor activity can be assessed by the open field tests via measuring distance moved, number of rearing, and time duration in the center and immobility time. Here, we firstly found that propionate administration was able to reverse reduced distance traveled, number of rearing, and time spent at the center of the open field (p<0.05) caused by STZ treatments (p<0.05) (Fig. 8A, B and C). No statistical significance was presented on these three indexes between propionate and metformin groups. The immobility time was shown in Fig. 8D. Mice in the STZ group spent more immobility time than those in the control group (p<0.001). Nevertheless, propionate reversed this result (p<0.001). Notably, propionate-treated mice presented remarkably less immobility time than metformin-administrated mice (p<0.05). Therefore, the results from open field tests suggested that propionate could enhance the exploratory willingness of STZ-induced diabetic mice.

**Fig. 7.**
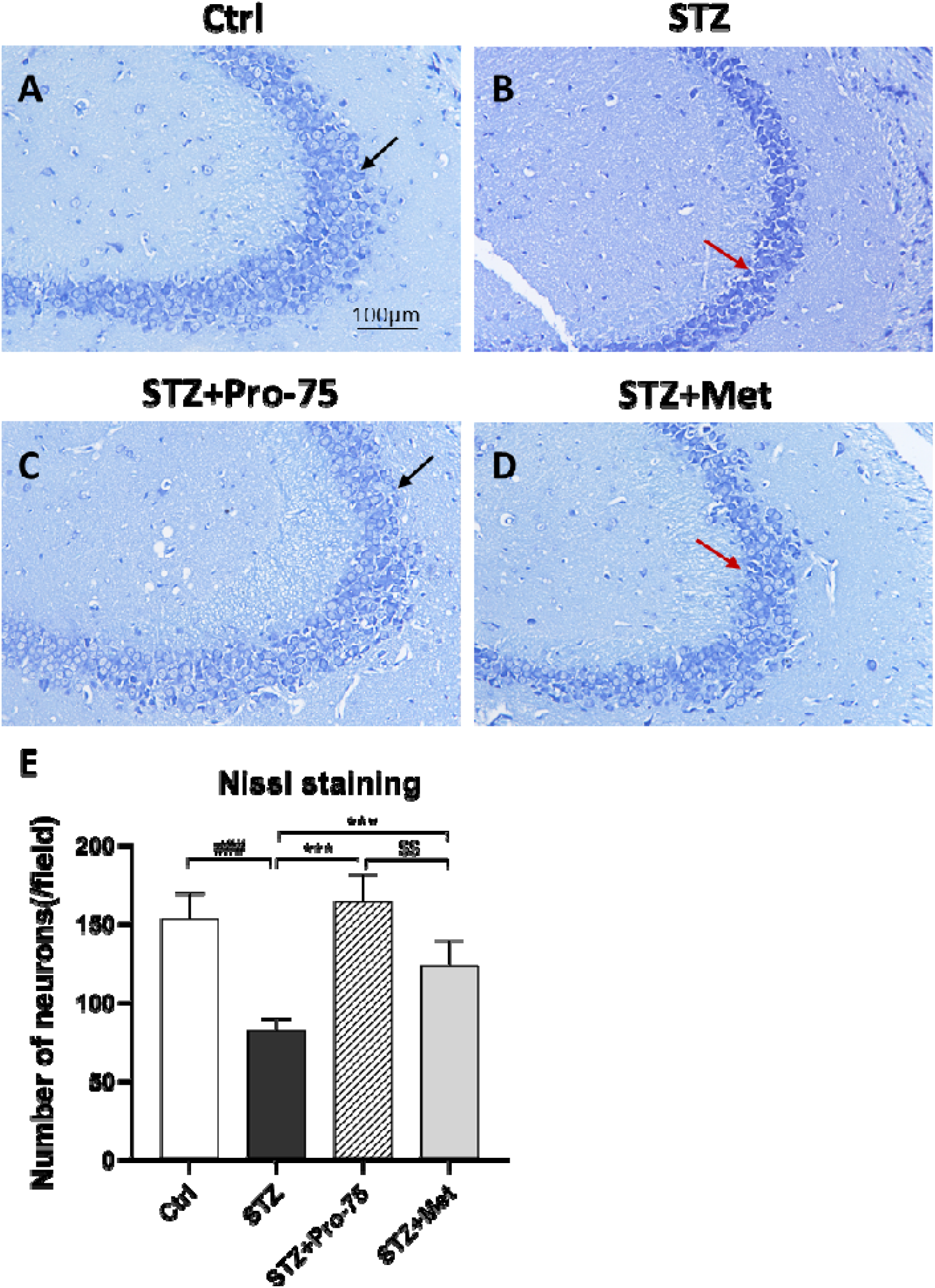
Cellular structures in the hippocampus are visible after Nissl staining. (A-D) Representation images of Ctrl, STZ, STZ + Pro, and STZ + Met, 36 days after IDDM induction. (E) Quantitative of neurons in the CA3. The red arrows indicate normal neurons, while the black arrows indicate necrosis neurons. Scale bar = 100 μm. Data were expressed as mean ± SD (N = 3). ###p < 0.001 vs. Ctrl group, ***p < 0.001 vs. STZ group, ^$$^p < 0.01 vs. STZ + Met group.

**Fig. 8.**
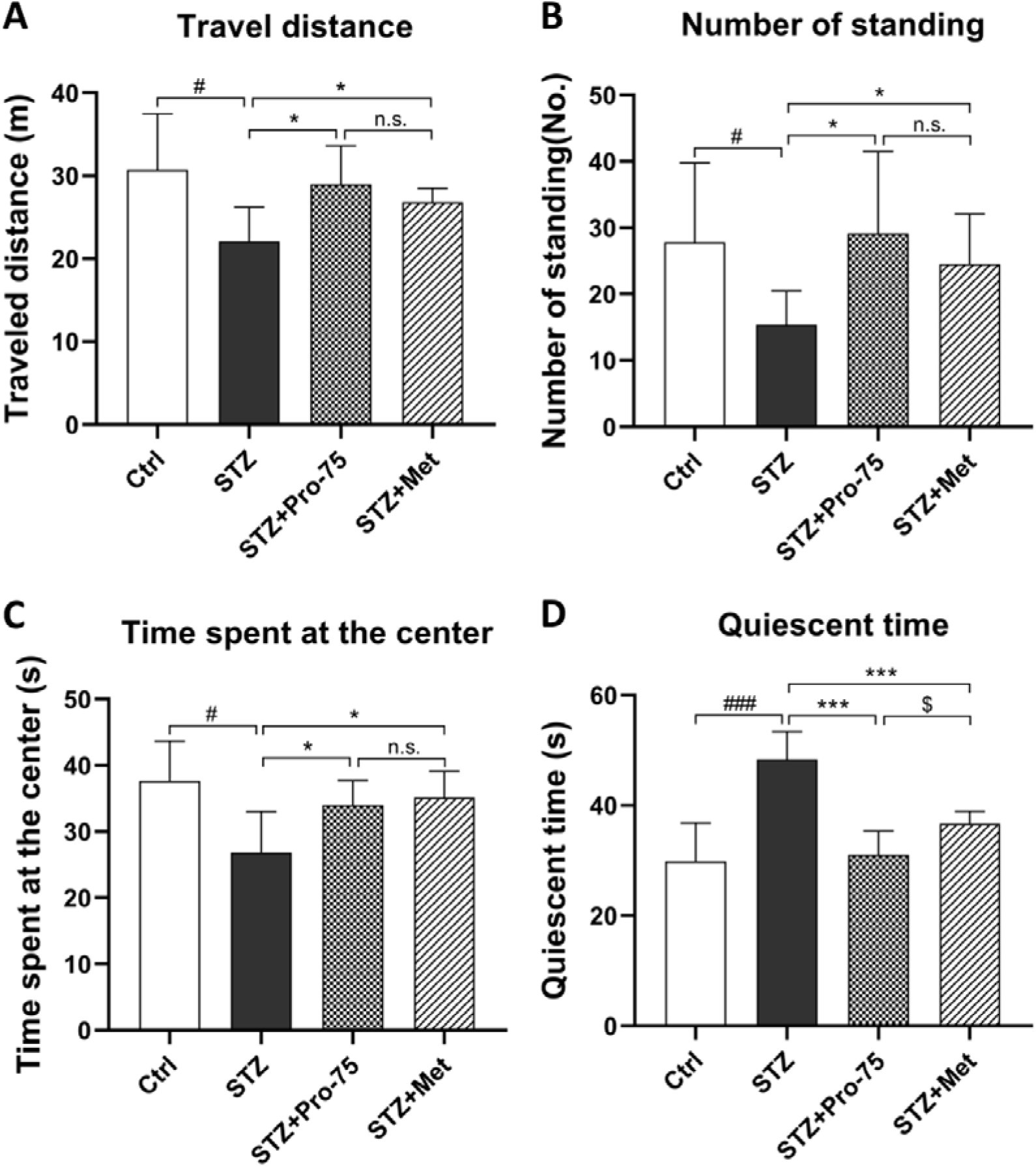
Animal behavior analysis in the open field test. In the OFT, the travel distance (A), number of standing (B), time spent at the center (C) and immobility time (D) were recorded and analyzed. Data were expressed as mean ± SD (N = 10). #p < 0.05 and ###p < 0.001 vs. Ctrl group, *p < 0.05 and ***p < 0.001 vs. STZ group, ^$^p < 0.05 vs. STZ + Met group, n.s., no significant difference.

### 3.8. Antidepressant effects of propionate treatment

Next, these results were further confined in an elevated plus-maze, which is wildly utilized to investigate the anxiety state of animals. As illustrated in Fig. 8, mice treated with STZ were less inclined to enter or remain in open arms (p<0.01 and p<0.05, respectively). However, propionate-treated or metformin-treated mice entered open arms more frequently than those in the STZ group (Fig. 9A, p<0.05). In addition, the retention time in open arms was significantly longer in the propionate-treated group than STZ model group (p<0.05). Interestingly, mice in the propionate group tarried for a longer period in open arms than mice in the metformin group (p<0.05). By contrast, metformin-treated mice showed no significant difference in time in open arms with those in the STZ group (Fig. 9B). Thus, the results in open arm tests indicated that propionate treatment attenuated anxiety in the STZ-induced IDDM model.

**Fig. 9.**
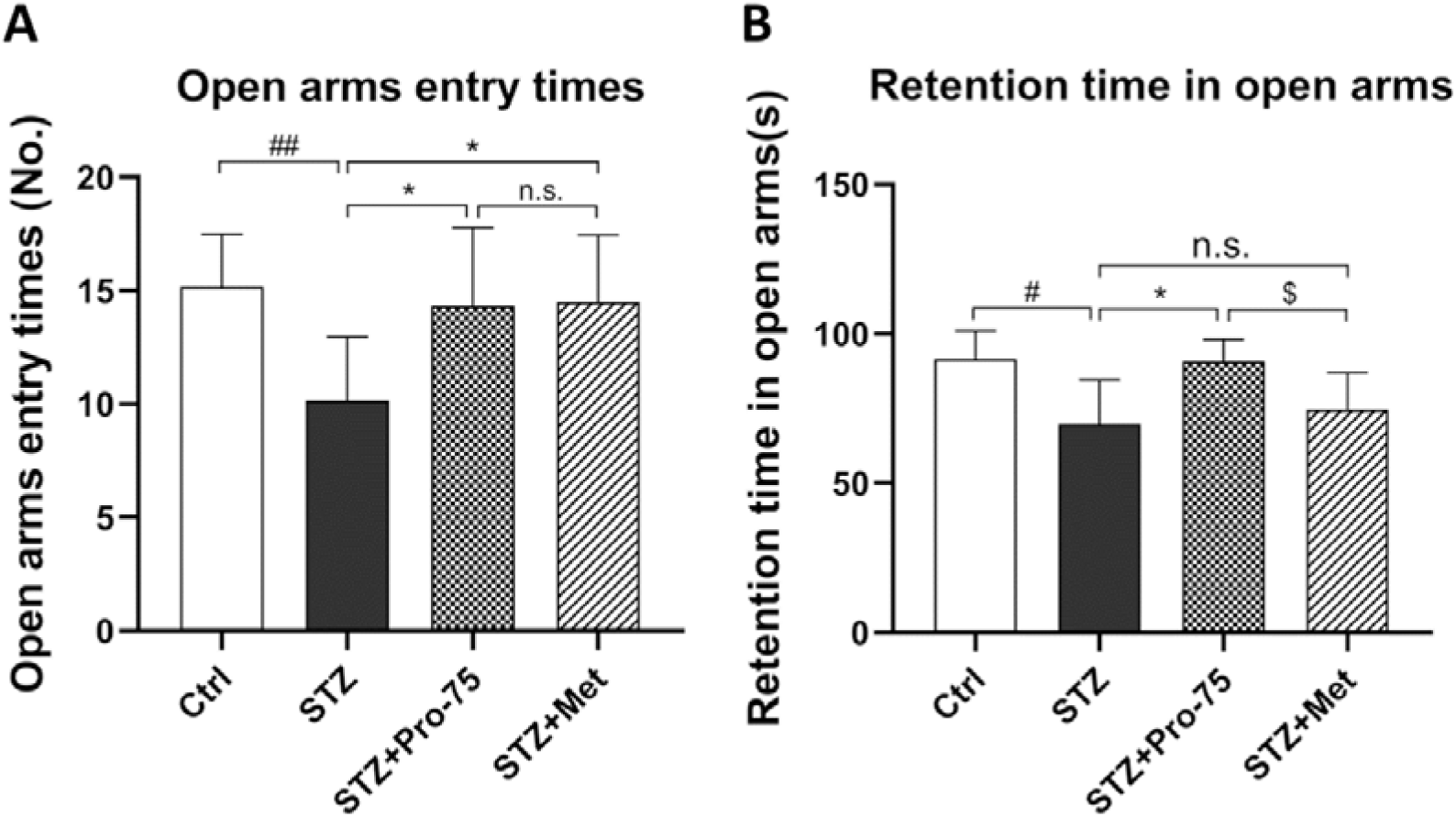
Results of elevated plus maze test. Open arms entry times (A) and retention time in open arms (B) were recorded and analyzed. Data were expressed as mean ± SD (N = 10). #p < 0.05 and ##p < 0.01 vs. Ctrl group, *p < 0.05 vs. STZ group, ^$^p < 0.05 vs. STZ + Met group, n.s., no significant difference.

Moreover, the effects of propionate on ameliorating mice’s depression-like behaviors were estimated in tail suspension tests and forced swimming tests. As shown in Fig. 10A, STZ-induced diabetic mice presented significantly longer immobility time when suspended by tails (p<0.05). Both propionate and metformin diminished the immobility time (p<0.05 and p<0.01, respectively). STZ exposure reduced the intention of tail climbs (Fig. 10B, p<0.001), while propionate or metformin treatments significantly reversed these tendencies (p<0.001 and p<0.05, respectively). Interestingly, propionate showed a better effect in increasing tail climbing behaviors than metformin (p<0.05). Furthermore, mice in the STZ-treated model group showed a remarkable increase in immobility time than mice in the control group in forced swimming tests (Fig. 10C, p<0.05), which was significantly reduced with the presence of propionate or metformin administrations (p<0.05), although the immobility time in STZ + Pro group was still longer than STZ + Met group (p<0.05). Therefore, the results implied that the administration of propionate could ameliorate the depression-related status of mice induced by STZ induction.

**Fig. 10.**
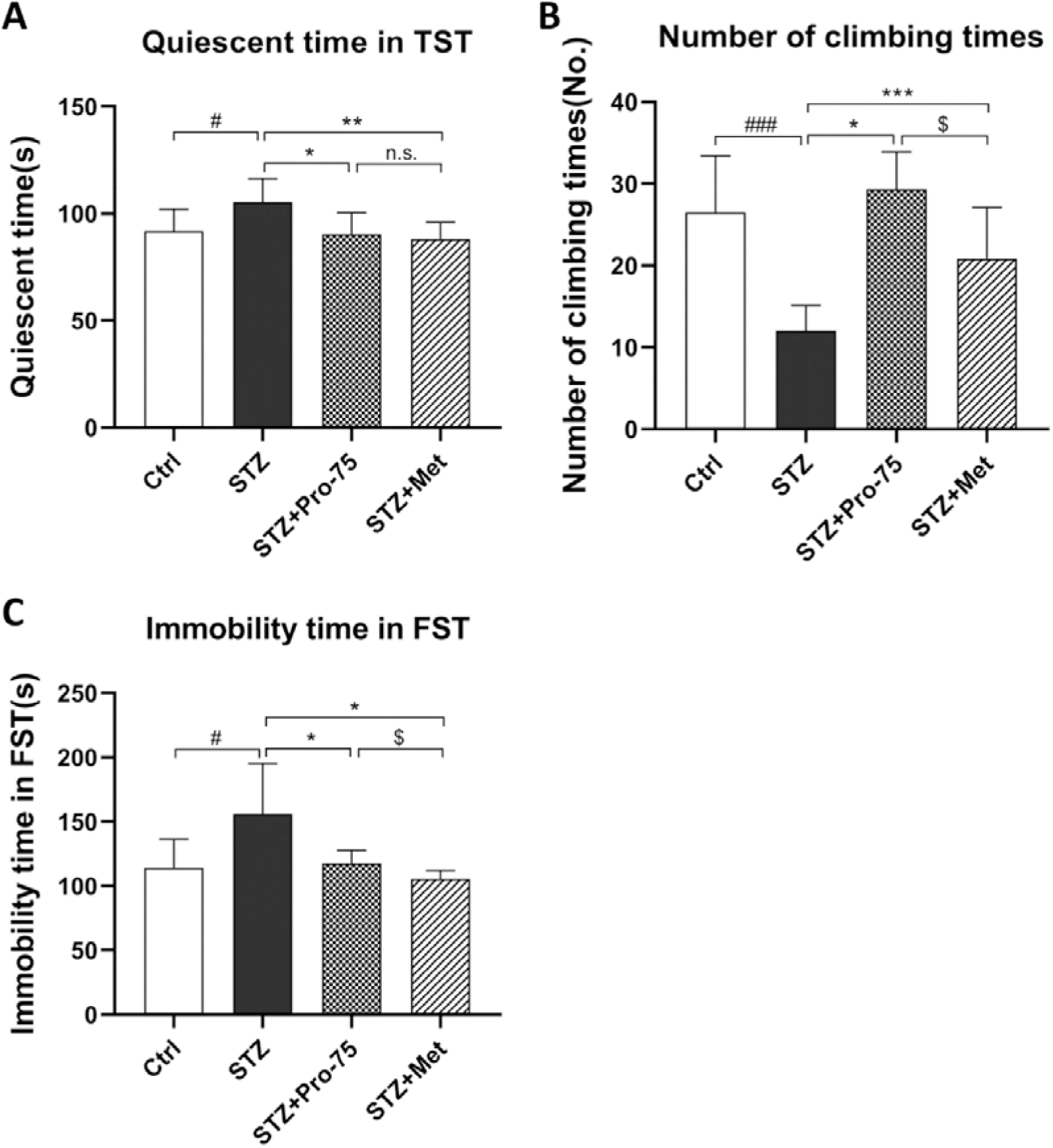
The influence of propionate on STZ-induced depression-like behavior evaluated by tail suspension test (TST) and force swimming test (FST). Quiescent time in TST (A), number of climbing times (B), and immobility time in FST (C) were measured and analyzed. Data were expressed as mean ± SD (N = 10). #p < 0.05 and ###p < 0.001 vs. Ctrl group, *p < 0.05, **p < 0.01, ***p < 0.001 vs. STZ group, ^$^p < 0.05 vs. STZ + Met group, n.s., no significant difference.

### 3.9. Improvement of learning and memory by propionate

Spatial memory functions of mice were commonly evaluated by water maze tests. As shown in figure 11A and Table 2, we found a significantly longer escape latency of the STZ group than the control group on day 2 and day 3-4 (p<0.05 and p<0.001, respectively), although not on day 1. Compared with the STZ model group, propionate treatment significantly reduced the escape latency of mice on day 2-4 (p<0.01 on day 2 and 3; p<0.001 on day 4). Metformin also decreased the latency on day 2 (p<0.001) and day 4 (p<0.05) but the propionate group showed a significantly shorter latency time than the metformin group in the last two days (p<0.05 for day 3 and p<0.01 for day 4). In summary, propionate and metformin could positively alter spatial learning of diabetic mice, while propionate was found to have better efficiency.

**Table 2.**
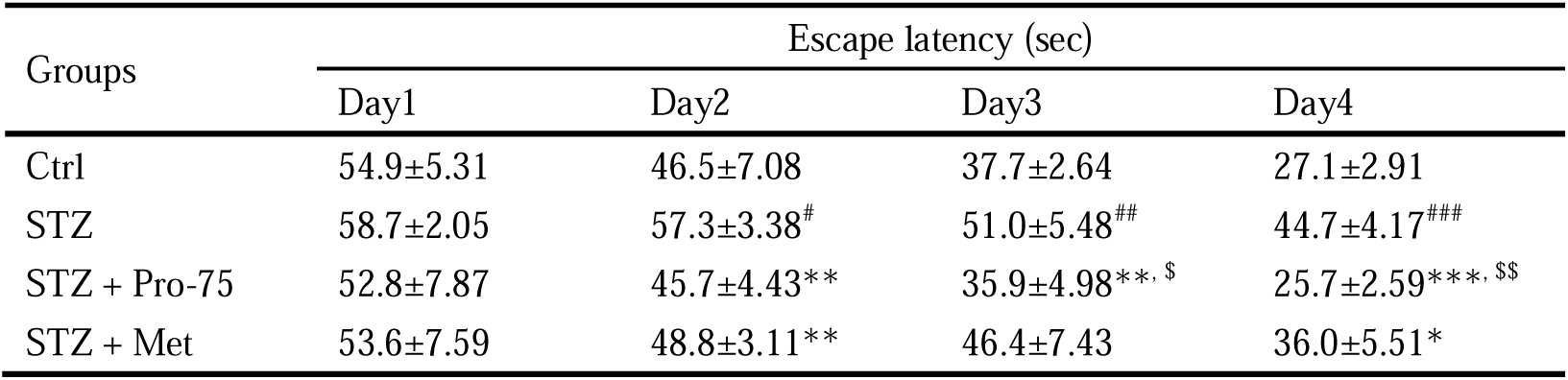
The exact escape latency in morris water maze values in seconds. Data was expressed as mean ± SD (N = 10). #p < 0.05, ##p < 0.01, and ###p < 0.001 vs. Ctrl group, *p < 0.05, **p < 0.01, and ***p < 0.001 vs. STZ group, ^$^p < 0.05 and ^$$^p < 0.01 vs. STZ + Met group. The two-way ANOVA for escape latency indicated a significant group-by-day interaction.

**Fig. 11.**
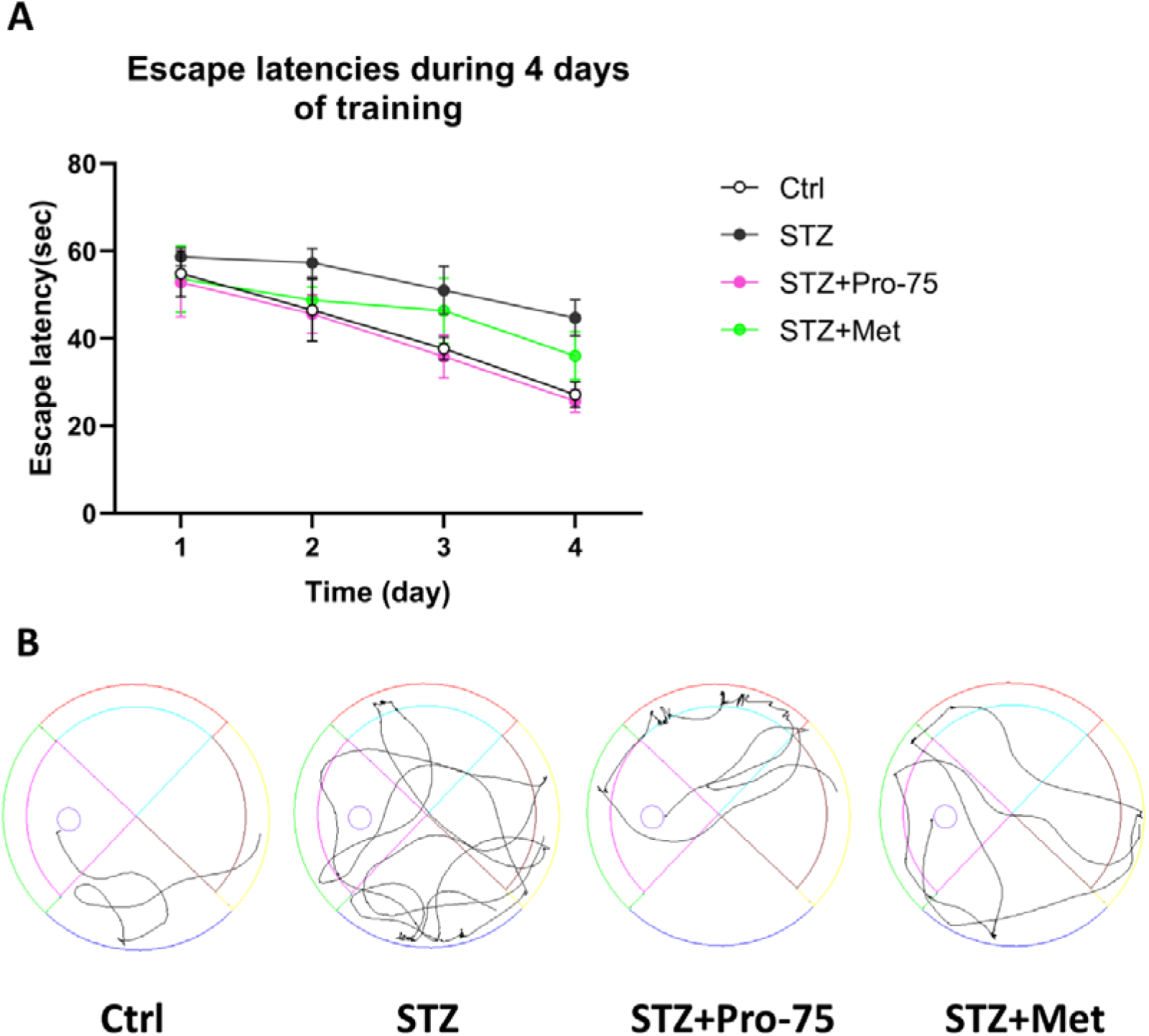
Escape latencies are shown as the mean value of each trail session for four sequential days (A). Swimming tracks are recorded via a video tracking camera system (B). Data for the escape latencies were analyzed with a two-way analysis of variance (ANOVA). Data was expressed as mean ± SD (N = 10). #p < 0.05 vs. Ctrl group, *p < 0.05 and **p < 0.01 vs. STZ group, n.s., no significant difference.

The number of crossing into the former location of the platform was also recorded (Fig. 12A). The STZ treated group showed a significant decrease in crossing number in comparison with the control group (p<0.05). Both metformin treatment and propionate treatment significantly restored the crossing number (p<0.05). Another indicator, time spent in the target quadrant in probe task, was significantly lower in STZ treated group in comparison to the control group (p<0.01) (Fig. 12B). While both the propionate-treated mice and metformin-treated mice spent a significantly longer time in the target quadrant than the STZ group (p<0.001 and p<0.01, respectively). Besides, the time that propionate-treated mice spent was longer than the metformin-treated group (p<0.05). As for the swimming velocity, there had been a falling-off in STZ treated group (p<0.05) (Fig. 12C). In contrast, the propionate-treated mice were faster (p<0.05), while the metformin-treated mice showed no significant acceleration compared to the propionate pretreated mice.

**Fig. 12.**
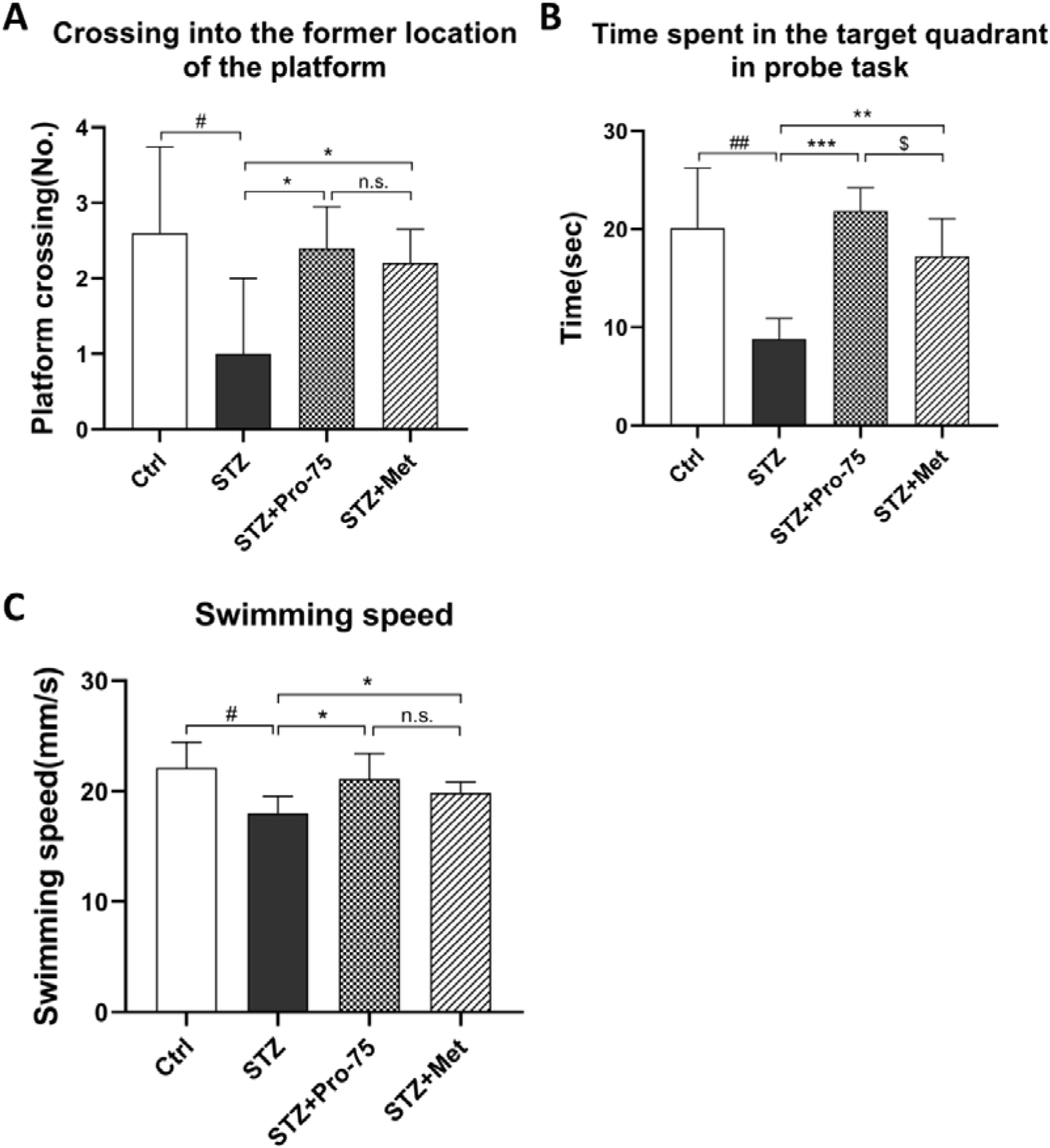
Influence of propionate on STZ-induced cognition and memory dysfunction measured by MWM test. Crossing into the former location of the platform (A), time spent in the target quadrant in probe task (B) and Swimming speed (C) were recorded and analyzed. Data were expressed as mean ± SD (N = 10). #p < 0.05 and ##p < 0.01 vs. Ctrl group, *p < 0.05, **p < 0.01, and ***p < 0.001 vs. STZ group, ^$^p < 0.05 vs. STZ + Met group, n.s., no significant difference.

**Fig. 13.**
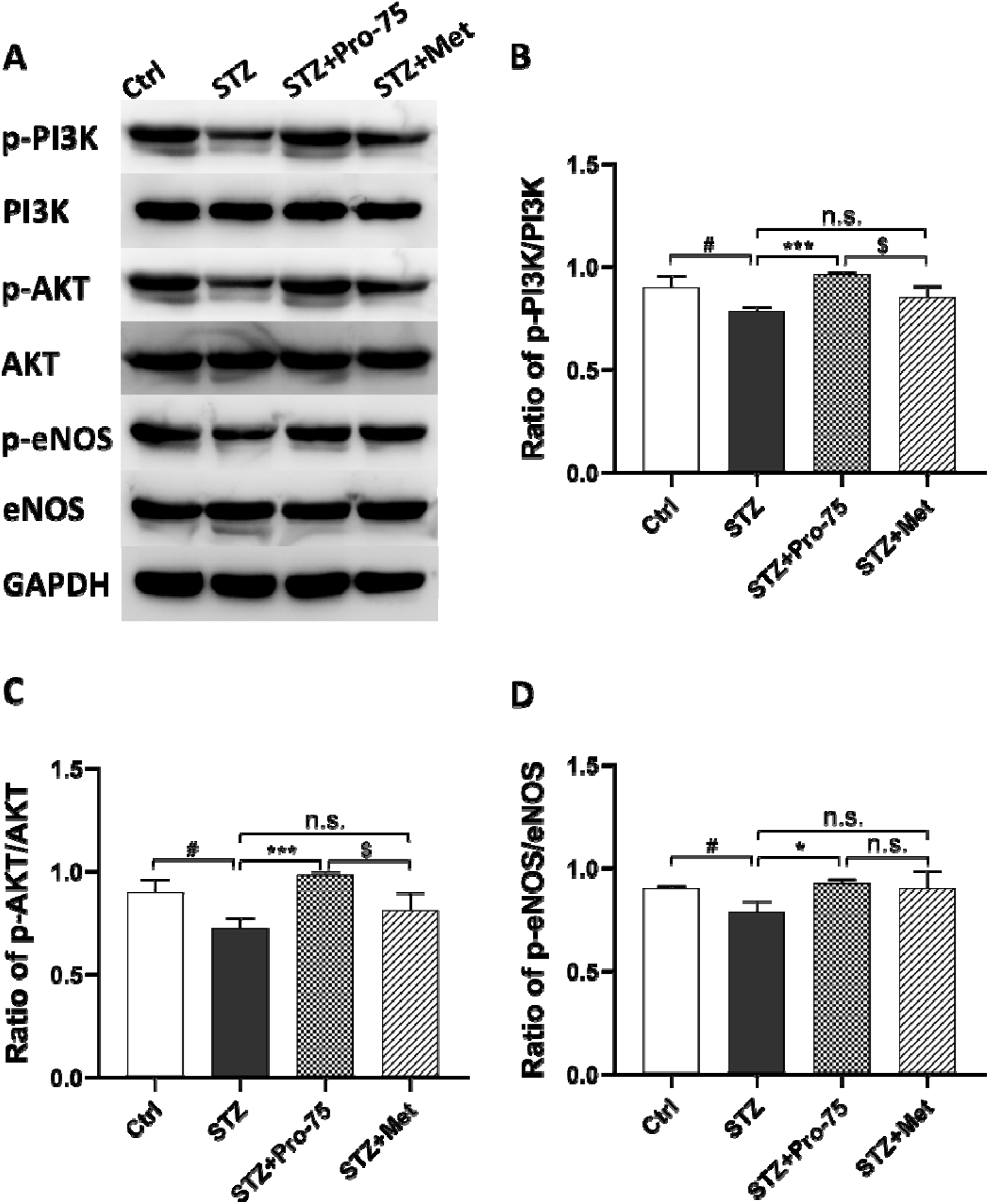
Phosphorylation of PI3K/AKT/eNOS signaling pathway proteins, as measured by western blotting. Representative western blot images (A) and the ratio of p-PI3K/PI3K (B), p-AKT/AKT (C), and p-eNOS/eNOS. GAPDH served as an internal control. Data are presented as the mean ± standard deviation (n=3 independent experiments). #p < 0.05 vs. Ctrl group, *p < 0.05 and ***p < 0.001 vs. STZ group, ^$^p < 0.05 vs. STZ + Met group, n.s., no significant difference.

### 3.10. Potent activation of the PI3K/Akt/eNOS pathway by propionate

Propionate activates the PI3K/Akt/eNOS pathway in the hippocampus of diabetic mice. Western blotting was implemented in this study for the purpose of identifying the relative expression of phosphorylated PI3K, Akt, and eNOS. From the panel, the protein levels of PI3K, Akt and eNOS remained unaltered acrossing all 4 groups. While, the STZ significantly reduced the phosphorylation levels of PI3K, Akt and eNOS. Both metformin and propionate presented rescue effects on phosphorylation level of PI3K, Akt and eNOS comparing with the STZ group (p<0.001 for PI3K and Akt; p<0.05 for eNOS). Propionate had exserted stronger effects on restoring phosphorylation levels of PI3K and Akt from the comparison between the propionate group and metformin group (p<0.05).

## Discussion

Diabetes mellitus is the most common chronic metabolic syndrome is characterized by absolute or relative insulin deficiency. The major symptom is chronic hyperglycemia caused by disturbance of saccharides, lipid, and protein metabolism ^43^. According to an epidemiological survey in 2015, approximately 8% of the global population aged between 20 to 79 suffers from diabetes, and the proportion is anticipated to rise to 9.9%. Abundant studies consistently suggest that diabetes mellitus (DM) affects the nervous system. Due to the prolonged life expectancy of DM patients, cognitive impairment and dementia have become new striking complications of DM ^44^. DM-induced cognitive deficits are also known as diabetic encephalopathy ^45^. Presently, many studies on diabetic encephalopathy worldwide focus on peripheral neuropathy, while the pathophysiological mechanisms of neuropathy caused by diabetic encephalopathy are less explored. However, with the significant increase in the life expectancy of DM patients, the prevalence of diabetic cognitive dysfunction is bound to increase. Thus, illuminating the pathogenesis of diabetic cognitive dysfunction and seeking effective protective or therapeutic strategies are urgently required.

This study contributes to this research field by discovering the therapeutic effect of propionate on IDDM, which has not been discussed previously. Firstly, we noticed that propionate intervention decreased blood glucose levels in STZ-induced type I diabetic mice, reversed their weight loss, increased their local cerebral blood flow, and indirectly improved their neural function. Secondly, propionate showed good antioxidant activity and could alleviate the loss of neurons in the hippocampus in IDDM mice. Moreover, we discovered that the neural protective function of propionate might be associated with the modulation of NO biosynthesis and improved blood supply to brain tissues. Furthermore, we found that the brain-protective function of propionate in STZ-induced IDDM mice was related or at least partly related to PI3K/Akt/eNOS signaling pathway.

STZ is a broad-spectrum antibiotic. It has properties of antibacterial, antitumor, and causing diabetes, and it has high-selective toxicity of pancreatic β-cells in experimental animals. Its activity in inducing diabetes may be through free radical damage to islet β-cells, which impairs the insulin-producing function of islet β-cells and leads to diabetes. Intraperitoneal injection of STZ has recently been applied in many diabetes-related studies to specifically destroy pancreatic β-cells, resulting in secretion disorder and persistent hyperglycemia without toxicity to brain cells ^46^. In this study, after a single intraperitoneal injection of STZ at 55 mg/kg to mice, their body weight was significantly decreased, and the fasting blood glucose level could reach up to or more than 20 mmol/L. Mice treated with STZ were generally in poor conditions, and some of them died before the terminal of the study.

Even in pediatric patients, metformin is the most commonly given treatment for hyperglycemia. ^47, 48^, It belongs to the biguanide family of drugs and works to lower blood sugar levels by reducing hepatic gluconeogenesis and enhancing insulin sensitivity. While metformin has been used to treat diabetes for more than 60 years, new evidence suggests that it could be used to treat a variety of other diseases, including cancer, cardiovascular disorders, neurodegenerative diseases ^49^ and more recently, intellectual disability-related neurodevelopmental diseases ^50, 51^. Metformin’s mechanism of action is not totally understood. Metformin’s activity is clearly focused on energy metabolism, and accumulating data shows that its therapeutic potential may be derived from its ability to target mitochondrial abnormalities. ^52^. In fact, chronic metformin administration enhances mitochondrial activity and antioxidant protection. Consistent with metformin’s propensity to cross the blood-brain barrier, these therapeutic effects have been observed in both the peripheral and central nervous systems. ^53^. Thus, metformin was used as a positive control drug in this study.

Insufficient insulin secretion leads to hyperglycemia. Persistent high blood glucose may cause thickening of capillary basement membrane and narrowing of vessel lumen ^54^. In combination with increased blood viscosity and slowed blood flow caused by metabolic disturbance of saccharides and lipids in diabetic patients, the structure changes of capillaries would decrease cerebral blood flow ^55^. Some research has proved that the decrease of cerebral blood flow could cause dysfunction of the brain in information processing, manipulating, and integrating ^56^. It would also impact cognitive response and processing ability, and eventually lead to impairment in learning and memory. In this study, cerebral blood flow was measured, besides the body weight and blood glucose level of mice. According to the results, in STZ-induced diabetic mice, their blood glucose level was markedly increased, and body weight was significantly decrease related to the control mice 36 days after STZ induction. As expected, results from cerebral blood flow studies also demonstrated obvious lower blood flow in high-glucose mice (36 days after STZ administration) than control mice. In contrast, after sodium propionate treatment, the blood glucose level of STZ-induced IDDM mice was significantly reduced. The sodium propionate exhibited the best effect on controlling blood glucose with a dose of 75 mg/kg. The weight gain of mice in the sodium propionate group was improved compared with the simple model group. Moreover, the cerebral blood flow of the diabetic mice was significantly increased after 36 days compared with the model mice, as showed by results from cerebral blood flow measurement.

Studies have found that long-term hyperglycemia is closely related to diabetic neurological complications, and diabetic models have exhibited varying degrees of behavioral and cognitive dysfunction ^57^. In this study, the behavior of experimental mice was evaluated by animal ethology methods, in order to study the improvement brought by propionate in neural function. The open-field test was used to evaluate the activity and exploration desire of mice, the tail suspension test and forced swimming test was used to evaluate the depression-like behaviors, and the water maze test was used to detect spatial learning and memory ability of mice. Experimental results showed that long-term hyperglycemia-induced significant cognitive and behavioral impairments, which are concretely showed in depression, a decline of learning and memory, and the results were consistent with that of previous studies about cognitive and behavioral impairments in diabetic models ^58^.

It has been reported that high glucose leads to excessive production of reactive oxygen species (ROS) in vascular endothelial cells ^59^. Excessive ROS formation affects the hypoxia signaling pathway and further aggravates vascular endothelial disorders ^60^. It causes damage to lipids, proteins and mitochondrial DNA on the mitochondrial intima of neurons, resulting in intracellular energy failures and diabetic brain injury. At the same time, oxidative stress will result in metabolic disorders. The results of this study showed that the hippocampus CA3 area of the experimental diabetes model mice was significantly damaged, and the neuron loss was severe. The SOD and CAT activities in the brain tissue were significantly decreased, while the MDA content increased. The preventive intervention of propionate can significantly ameliorate the morphological changes of brain tissue and reduce the loss of neurons in the hippocampus, and antioxidant effects were also significantly enhanced. These findings indicate that propionate has a significant anti-oxidative function and can relieve hyperglycemia-induced brain damage in mice while promoting the regeneration of neurons.

Studies have shown that vascular endothelial cells can secrete a variety of vasoactive substances, including vasodilator factor NO. As a previous study suggested, NO synthesis is inhibited due to impaired vascular endothelial function in diabetic patients. Hyperglycemic environment leads to a reduction in endothelial cell function and reduction in the bioavailability of NO. In addition, high blood sugar culture conditions will accelerate the senescence of vascular endothelial cells, leading to impaired proliferation ^61^. This study found that the NO level of model mice was significantly reduced, and propionate treatment significantly increased the NO level of mice. It is suggested that the therapeutic effect of propionate may be related to the regulation of NO synthesis and secretion. Despite the fact that NO is a byproduct of oxidative stress, it was found to have a protective impact on the brain in this study. The findings of the experiments revealed that the primary function of NO is to widen blood vessels, protect vascular endothelium, and boost blood supply to the brain. It may be observed that the preventive impact of NO outweighs its oxidative harm. Other researchers have found that inhibition of the PI3K/Akt/eNOS signaling pathway can engender vascular endothelial cell dysfunction, and a decrease in NO synthesis is observed when eNOS activity decreases since the eNOS is mainly activated by the PI3K/Akt signaling pathway. In this study, propionate treatment was shown to significantly upregulate the expression levels of PI3K/Akt/eNOS signaling pathway-related proteins, suggesting that propionate’s brain protection function in STZ-induced type I diabetic mice is associated with PI3K/Akt/eNOS signaling pathway activation.

Our findings demonstrated that propionate (75 mg/kg) has a beneficial effect on the injured brains of mice. As a result, approximately 0.5 g propionate should be administered to a 60 kg individual via dose conversion from mice to human. Propionate levels in bread range between 1.0 and 2.5 g per kilogram. We believe that the recommended dosage of propionate is easily achieved from food supply. Supplements in the diet, such as propionate, appear to be promise for safeguarding the brain against harm in IDDM. However, given the difference between humans and mice, further rigorous studies examining the safety and efficacy of such high doses of propionate in different animal models of epilepsy are required prior to initiating clinical trials.

## Conclusion

Taken together, strong evidence has been provided that propionate acts to ameliorate cerebral microcirculation to improve the cognitive abilities in diabetic mice, which may be attributed to its anti-diabetic, antioxidant and modulated by PI3K/Akt/eNOS pathway. The results of this study emphasize the importance of propionate on NO release, cerebral circulation, oxidative stress, and PI3K/Akt/eNOS pathway against high glucose insult.

## Abbreviations

IDDM: insulin-dependent diabetes mellitus
DM: diabetes mellitus
STZ: streptozocin
Met: metformin
Pro: propionate
NO: nitric oxide
PYY: peptide YY;
GLP-1: glucagon like peptide-1;
ROS: reactive oxygen species;
SOD: superoxide dismutase
CAT: catalase;
MDA: malondialdehyde
ROI: region-of-interest
LCBF: local cerebral blood flow
ELISA: enzyme-linked immunosorbent assay
PI3K: phosphoinositide 3-kinase
Akt: serine-threonine protein kinase
eNOS: endothelial nitric oxide synthase
PDK1: 3-inositol phosphate-dependent protein kinase-1
TST: tail suspension test
FST: force swimming test
MWM: Morris water maze
FBG: fasting blood glucose
IPGTT: intraperitoneal glucose tolerance test
SDS-PAGE: sodium dodecyl sulfate-polyacrylamide gel electrophoresis
PVDF: polyvinylidene fluoride
Ctrl: control
ANOVA: analysis of variance
SD: standard deviation
AUC: area under the curve

## Funding source

This study did not get any funding from funding agencies

## Conflict of interest

The authors acknowledge no conflict of interest pertaining to the publication of the manuscript.

## Acknowledgements

Thanks to Shiqi Fan and Jinjin Li for their help in preparing samples and in measuring biochemical indices.

## Author contributions

Yahong Cheng: Conceptualization, Writing-original draft and Writing-review & editing.

Yuan Jiang and Xinying Bai: Data curation and Methodology.

Qin Wu and Jiajun Dong: Conceptualization, Software, Data curation and analysis, Methodology,

Writing-original draft, drawing graphic abstract, and revising the manuscript.

Gaofeng Jiang: Supervision, Validation and Methodology.

## Notes

### Competing Interest Statement

The authors have declared no competing interest.

